# Human CD8^+^ T cells exhibit a shared antigen threshold for different effector responses

**DOI:** 10.1101/2020.04.24.059766

**Authors:** Enas Abu-Shah, Nicola Trendel, Philipp Kruger, John Nguyen, Johannes Pettmann, Mikhail Kutuzov, Omer Dushek

**Affiliations:** Sir William Dunn School of Pathology; Kennedy Institute of Rheumatology University of Oxford

**Author notes:** Equal contribution (Enas Abu-Shah, Nicola Trendel). **Corresponding author:** Omer Dushek, Sir William Dunn School of Pathology, University of Oxford, South Parks Road, Oxford, OX1 3RE, United Kingdom, E.

## Abstract

T cells recognising cognate pMHC antigens become activated to elicit a myriad of cellular responses, such as target cell killing and the secretion of different cytokines, that collectively contribute to adaptive immunity. These effector responses have been hypothesised to exhibit different antigen dose and affinity thresholds, suggesting that pathogen-specific information may be encoded within the nature of the antigen. Here, using systematic experiments in a reductionist system, where primary human CD8^+^ T cell blasts are stimulated by recombinant pMHC antigen alone, we show that different inflammatory cytokines have comparable antigen dose thresholds across a 25,000-fold variation in affinity. Although co-stimulation by CD28, CD2, and CD27 increased cytokine production in this system, the antigen threshold remained comparable across different cytokines. When using primary human memory CD8^+^ T cells responding to autologous antigen presenting cells equivalent thresholds were also observed for cytokine production and killing. These findings imply a simple phenotypic model of TCR signalling where multiple T cell responses share a common rate-limiting threshold and a conceptually simple model of antigen recognition, where the chance factor of antigen dose and affinity do not provide any additional response-specific information.

## Introduction

The activation of T cells is critical for immune responses that target infectious agents and cancers. T cells recognise antigen in the form of peptides presented on major-histocompatibility complex (pMHC) on professional antigen presenting cells and, in the case of CD8^+^ T cells, also directly on abnormal, infected or cancerous cells. To eradicate abnormal cells, they initiate a spectrum of effector responses such as direct target cell killing and production of a myriad of cytokines (1). Cytokines are important effectors that mediate cellular communication to promote inflammation (e.g. IFNγ and TNFα), proliferation (e.g IL-2), recruitment of other immune cells (e.g. MIP1β), interference with viral replication (e.g. IFNγ), along with many other functions (2–4). Given their critical function in cellular communication, it has been postulated that different cytokines exhibit a different antigen threshold for production (Fig. 1A).

**Figure 1:**
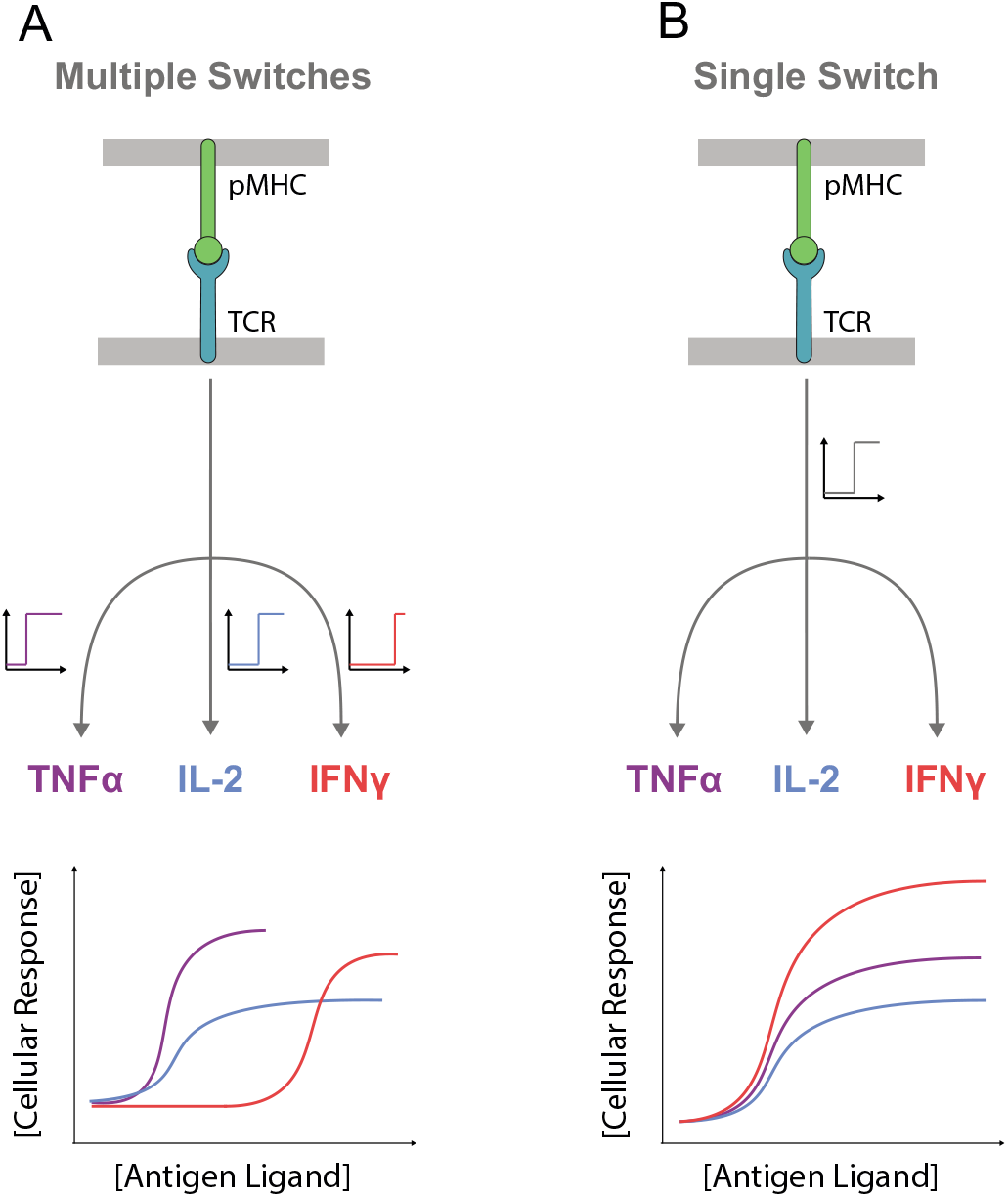
Phenotypic models for pMHC antigen induced digital TCR signalling leading to multiple T cell effector responses. A) Schematic of TCR signalling showing different rate-limiting digital switches controlling different cytokines. The antigen threshold for producing each cytokine can vary if the digital switch has a different threshold with respect to TCR signalling. B) Schematic of TCR signalling showing a common rate-limiting digital switch controlling different cytokines. The antigen threshold is identical because all cytokines are produced when the switch is “on”. In both models, the amount of cytokine produced can be regulated differently for each cytokine downstream of the switch.

Previous studies presented conditions where different T cell effector responses appeared to have different pMHC antigen thresholds (5, 6), and are differentially regulated by pMHC affinity and co-stimulatory molecules (7, 8). Moreover, different T cell clones exhibit a different hierarchical organisation of thresholds (9), implying that they have differential wiring of their signalling machinery. Those suggestions have strong implications to the immune response whereby T cells would be able to infer specific information from antigens generated by different pathogens. These studies often relied on peptide-pulsed antigen-presenting-cells (APCs), which although providing a physiological stimulus, have the caveat that the the antigen concentration and stability over time are difficult to control. This may produce apparently different thresholds if the kinetics of each response differ. Moreover, variation in the expression of ligands for co-signalling receptors on T cells can exist over time and differ between experiments. To our knowledge, systematic analyses controlling for these factors have yet to be performed.

T cell responses can exhibit a hierarchy in the antigen threshold dose and/or affinity if different effector responses exhibit different threshold sensitivities to TCR signalling. Given that TCR signalling is thought to be digital on the single-cell level (10, 11), one mechanism to produce different antigen thresholds would be to invoke a different rate-limiting switch for each response that has a different sensitivity to TCR signalling (Fig. 1A). On the other hand, if different responses shared a common rate-limiting switch then different responses would share a comparable antigen threshold (Fig. 1B). In the latter model, antigen affinity can control the threshold antigen concentration comparably for all responses. In both models, the production of cytokine can be regulated downstream of the switch so that the temporal kinetics and magnitude of the response (e.g. the amount of cytokine) can be different for different cytokines.

Here, we systematically stimulate primary human CD8^+^ T cell blasts in a reductionist system that allows for the precise control of pMHC antigen dose and affinity. We find that although antigen affinity controls the antigen dose threshold for inducing cytokine production, the threshold is comparable for different cytokines across a wide range of affinities. By incorporating ligands to CD28, CD2, and CD27, we show that although they increase cytokine production, they do so similarly for different cytokines so that the threshold remains comparable. Finally, we reproduce these findings in a recently described experimental system (12) that allows for the study of quiescent primary memory CD8^+^ T cells responding to autologous monocyte-derived dendritic cells. The work suggests a conceptually simpler phenotypic model for TCR signalling with implications for the role of antigen concentration and affinity in mediating T cell responses.

## Results

### Different cytokines exhibit comparable antigen dose thresholds over a wide range of affinity

To investigate the antigen threshold required to elicit different effector cytokines, we first utilised a reductionist system to exclude any contribution from extrinsic factors such as pMHC stability and variation in co-stimulatory ligands on APCs. In this system, primary human CD8^+^ T cell blasts that have been transduced to express the affinity enhanced 1G4 TCR (c58c61) (13, 14) were stimulated by plate-immobilised recombinant pMHC (15–18). The use of the c58c61 TCR allowed us to explore cytokine thresholds when T cells are stimulated by a panel of 8 pMHCs that span >10-million fold variation in affinity from supra-physiological therapeutic affinities (pM) to physiological affinities (μM) (Fig. S1) (17). After 8 hours of interacting with plates coated with different concentrations of the different affinity antigens, the production of TNFα, IFNγ, and IL-2 was quantified in the supernatants (Fig. 2A).

To compare the antigen threshold of each cytokine, we first determined the maximum efficacy (E_max_, the maximum response across all antigen doses) and the antigen potency (E_50_, the threshold concentration of antigen producing 50% of E_max_) by directly fitting the dose-response curves (see Methods). We then plotted representative experiments on the same graph either directly or normalised by the E_max_ of each cytokine to clearly identify whether any differences in antigen threshold were present (Fig. 2C, Fig. S2). This analysis shows that although a different amount of each cytokine is detected, the antigen threshold for inducing these cytokines is comparable and within the resolution of our 2-fold dilutions. This conclusion is reflected in the EC_50_ values for the 9 repeated experiments showing that no significant difference can be detected (Fig. 2C, Fig. S2, right two panels).

**Figure 2:**
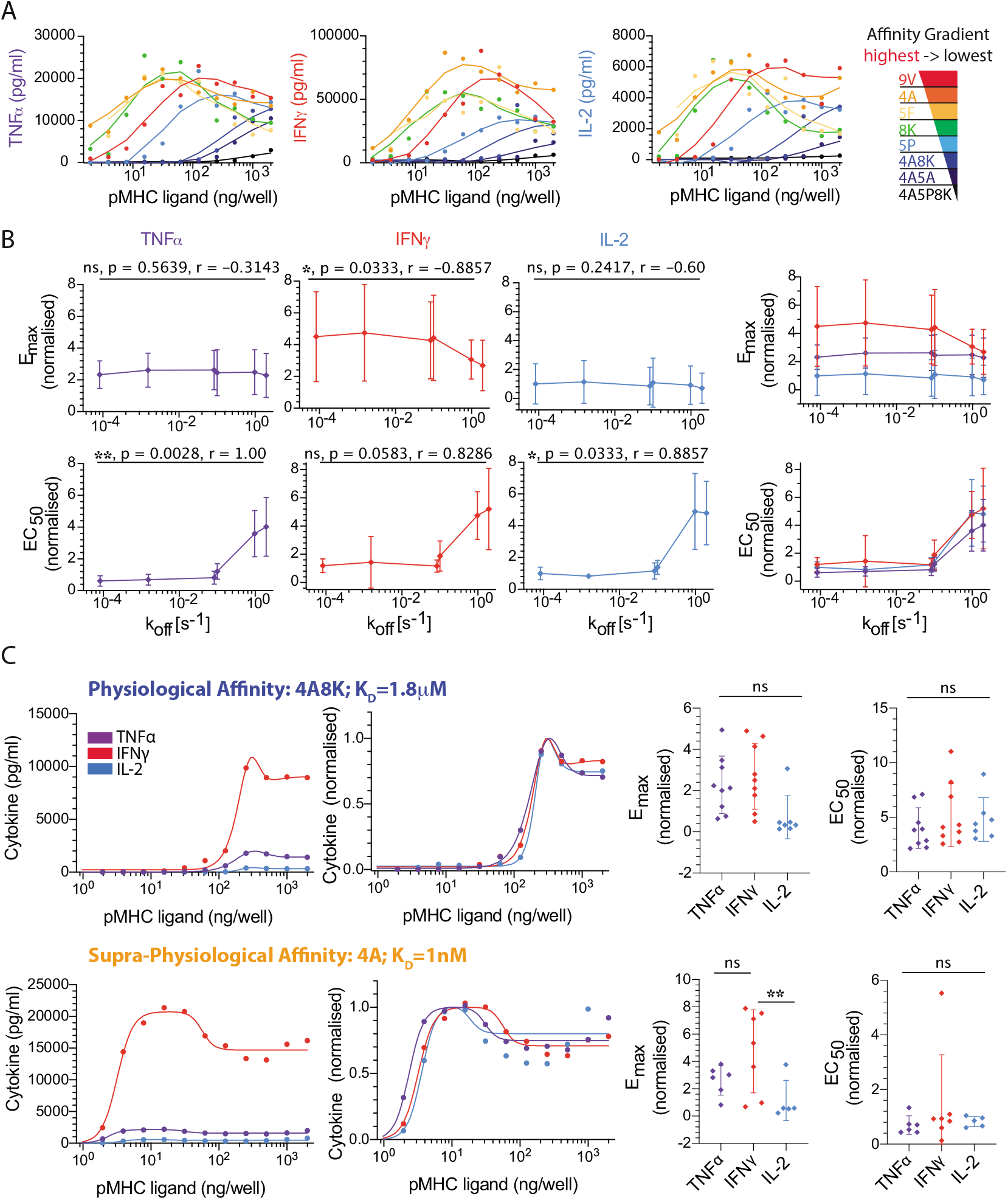
Different cytokines exhibit a comparable antigen dose threshold over a large variation in antigen affinity. A) Representative data showing supernatant TNFα, IFNγ, and IL-2 over pMHC dose for different pMHC affinities (colours). B) Fitted E_max_ (top row) and EC_50_ (bottom row) for each cytokine individually (left three columns) or overlaid (right column) over the TCR/pMHC *k*_off_ measured at 37°C (Fig. S1). Reliable estimates were not possible for the two lowest affinity pMHCs (4A5A, 4A5P8K) and they were omitted for quantitative analysis. C) Representative overlay of TNFα, IFNγ, and IL-2 directly (first column) or normalised by their respective E_max_ value (second column) for a physiological (top row) and supra-physiological (bottom row) affinity pMHC. Dot plots for E_max_ (third column) and EC_50_ (fourth column) show the data across 9 independent repeats with different donors. This analysis highlights that, within the resolution of our 2-fold dilutions in pMHC dose, no significant difference is observed between the EC_50_ threshold for different cytokines. ANOVA corrected for multiple comparisons by Tukey’s test (**; p=0.002). The normalised dose-response curves for all 9 repeats is displayed in Fig. S3. Error bars are SD of mean. Normalisation of experimental data is described in Materials and Methods. Solid lines in representative datasets are the fits used to extract E_max_ and EC_50_.

Although we focused on TNFα, IFNγ, and IL-2, we found that the same threshold applied for MIP-1β (Fig. S3). Therefore, different cytokines exhibit a comparable antigen dose threshold across a wide variation in antigen affinity (~25,000-fold, 9V to 4A8K) when T cell activation is mediated exclusively by pMHC through the TCR.

### Co-stimulation increases cytokine production but maintains a comparable antigen dose threshold for different cytokines

Given that we observed a comparable antigen dose threshold for different cytokines when stimulating T cells with pMHC antigen alone, we hypothesised that differences between cytokines may emerge when providing T cell co-stimulation. This hypothesis is motivated by the fact that in previous reports showing differential cytokine thresholds, antigen was expressed on the surface of antigen presenting cells in the context of ligands to various co-stimulatory receptors on T cells. To directly test this hypothesis, we used our reductionist experimental system to co-present pMHC with a titration of three prominent CD8^+^ T cell co-stimulatory ligands: CD86 (ligating CD28; Fig. 3), CD58 (ligating CD2; Fig. 4) or CD70 (ligating CD27; Fig. 5).

**Figure 3:**
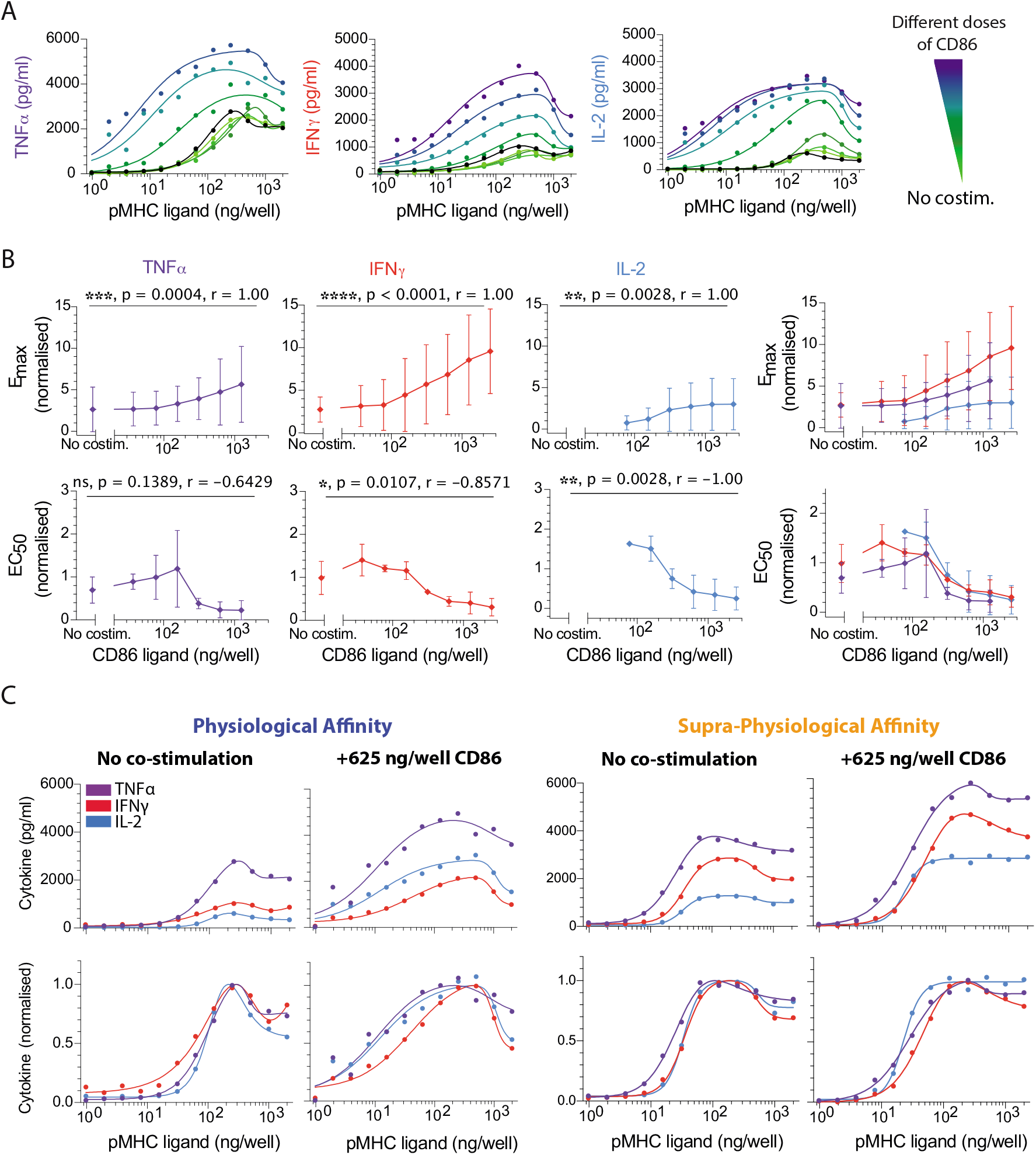
CD28 co-stimulation decreases the antigen threshold for cytokine production comparably for different cytokines. A) Representative data showing secretion of TNFα, IFNγ, and IL-2 over the pMHC dose (physiological affinity, 4A8K) when T cells were co-stimulated with different doses of CD86 (colours). Black solid line is without co-stimulation. B) Normalised E_max_ (top row) and EC_50_ (bottom row) for each cytokine over the CD86 dose confirms that co-stimulation can control both efficacy and potency, respectively. Overlay of E_max_ and EC_50_ for all cytokines (right most panels). C) Representative overlay of TNFα, IFNγ, and IL-2 directly (top row) or normalised (bottom row) for the indicated pMHC and co-stimulation condition. The antigen dose threshold for different cytokines is comparable irrespective of CD86 dose. For statistical comparison see Fig. S4. Error bars are SD of mean. Normalisation of experimental data is described in Materials and Methods. Solid lines in representative datasets are the fits used to extract E_max_ and EC_50_.

**Figure 4:**
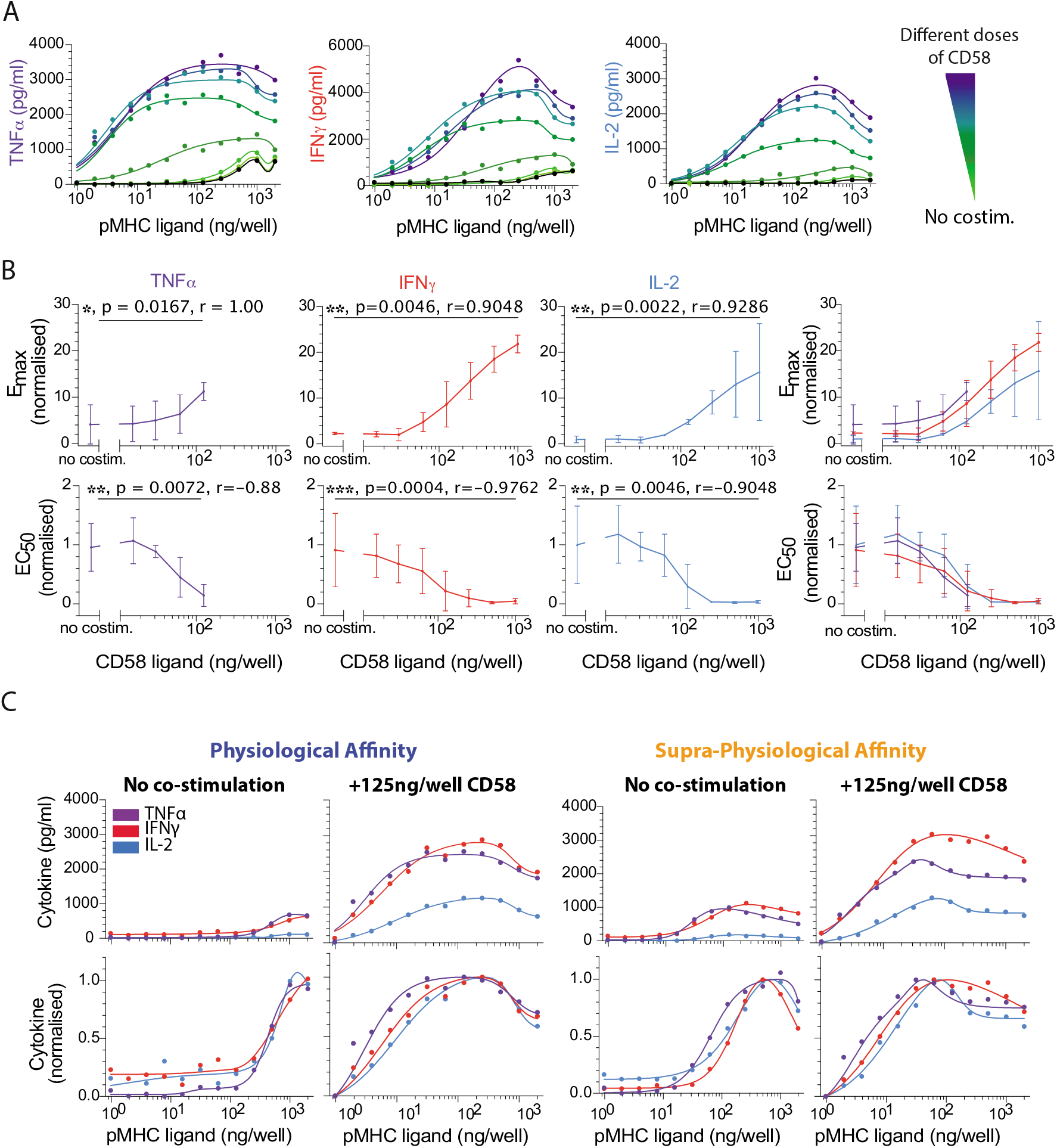
CD2 co-stimulation decreases the antigen threshold for cytokine production comparably for different cytokines. A) Representative data showing secretion of TNFα, IFNγ, and IL-2 over the pMHC dose (physiological affinity, 4A8K) when T cells were co-stimulated with different doses of CD58 (colours). Black solid line is without co-stimulation. B) Normalised E_max_ (top row) and EC_50_ (bottom row) for each cytokine over the CD58 dose confirms that co-stimulation can control both efficacy and potency, respectively. Overlay of E_max_ and EC_50_ for all cytokines (right most panels). C) Representative overlay of TNFα, IFNγ, and IL-2 directly (top row) or normalised (bottom row) for the indicated pMHC and co-stimulation condition. The antigen dose threshold for different cytokines is comparable irrespective of CD58 dose. For statistical comparison see Fig. S5. Error bars are SD of mean. Normalisation of experimental data is described in Materials and Methods. Solid lines in representative datasets are the fits used to extract E_max_ and EC_50_.

**Figure 5:**
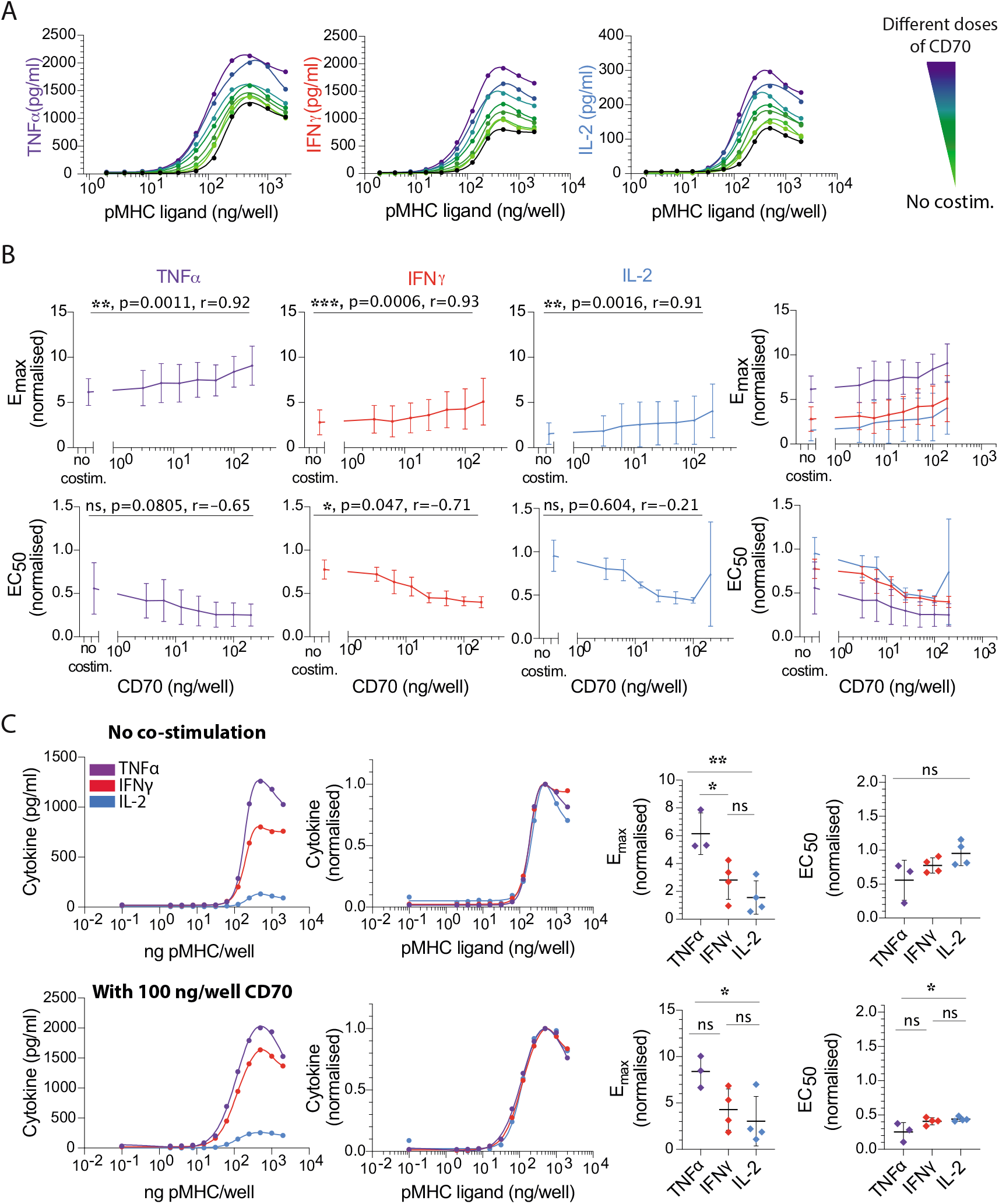
CD27 co-stimulation decreases the antigen threshold for cytokine production comparably for different cytokines. A) Representative data showing secretion of TNFα, IFNγ, and IL-2 over the pMHC dose (physiological affinity, 4A8K) when T cells were co-stimulated with different doses of trimeric CD70 (colours). Black solid line is without costimulation. B) Normalised E_max_ (top row) and EC_50_ (bottom row) for each cytokine over the CD70 dose confirms that co-stimulation can control both efficacy and potency, respectively albeit to a lower extent compared to CD28 and CD2. Overlay of E_max_ and EC_50_ for all cytokines (right most panels). C) Representative overlay of TNFα, IFNγ and IL-2 directly (first column) or normalised (second column) without co-stimulation (top row) or with a single dose of CD70 co-stimulation (bottom row). The antigen dose threshold for different cytokines is comparable under both conditions. This conclusion is reflected in the dot plots of E_max_ and EC_50_ showing no significant or up to 2-fold differences between the cytokines in the absence (top row) or presence (bottom row) of CD70. Error bars are SD of mean. Normalisation of experimental data is described in Materials and Methods. Solid lines in representative datasets are the fits used to extract E_max_ and EC_50_. ANOVA corrected for multiple comparisons by Tukey’s test (*,p=0.03; **, p=0.0052).

The ligand CD86 binds the co-stimulatory receptor CD28 (19). As expected, a titration of CD86 increased the amount of cytokine detected in the supernatant and reduced the antigen dose threshold required to detect cytokine (Fig. 3A,B). However, when we directly compared thresholds for individual cytokines, we observed that all responded quantitatively similar to individual CD86 concentrations across the different pMHC affinities (Fig. 3C, Fig. S4).

The ligand CD58 binds the co-stimulatory receptor CD2, which is thought to enhance adhesion (20) and intracellular signalling (21, 22). We observed a large ~50-fold increase in potency (decrease in EC_50_) and a > 10-fold increase in efficacy (Fig. 4A,B). However, as with CD28 co-stimulation, the antigen dose threshold appeared comparable for different cytokines under all conditions (Fig. 4C). We did observe a moderate difference for the supra-physiological pMHC affinity with low concentration of CD58, whereby the EC_50_ value of TNFα was ~2-fold lower compared to IFNγ and IL-2 (Fig. S5). However, given that this was only apparent under some conditions and similar in magnitude to the resolution of our assay (2-fold pMHC dilutions), it is unclear whether it is biologically relevant (see also discussion).

The ligand CD70 forms trimers that induce trimerisation of the co-stimulatory receptor CD27, which is a member of the TNF family of co-stimulatory molecules (23). We observed that a titration of recombinant and trimeric CD70 exhibited increased cytokine production (Fig. 5A) with a modest impact on both the efficacy and potency compared to CD2 and CD28 ligation (Fig. 5B). However, as with CD28 and CD2 co-stimulation, the antigen dose threshold for different cytokines appeared equivalent under all conditions (Fig. 5C).

Taken together, this data suggests that T cell co-stimulation by 3 prominent receptors that span diverse families can control cytokine production efficacy and potency but are unable to control potency differently for each cytokines we investigated.

### Memory CD8^+^ T cells exhibit comparable cytokine thresholds in response to autologus antigen presenting cells

Given that we observed comparable cytokine thresholds using T cell blasts in a reductionist system and that limited data is available for human T cells, it was important to determine if these conclusions hold in a more physiological system. To this end, we directly isolated memory CD8^+^ T cells from healthy donors and used RNA electroporation to express the wild-type 1G4 TCR, and in parallel, we generated autologous monocyte-derived dendritic cells that we loaded with a titration of the 9V peptide (K_D_ = 7.2 *μ*M (15)) before mixing them with T cells for either 6 or 24 hours (Fig. 6A). This experimental system has recently been described in detail (12). We focused on memory CD8^+^ T cells because naive T cells do not produce cytokines on our experimental timescales (24). Consistent with our previous results, we observed a comparable antigen threshold for all cytokines (Fig. 6B,C). We were also able to assess the ability of memory cells to kill target cells by measuring the release of LDH into the supernatant finding an antigen threshold that was comparable to the cytokine threshold.

**Figure 6:**
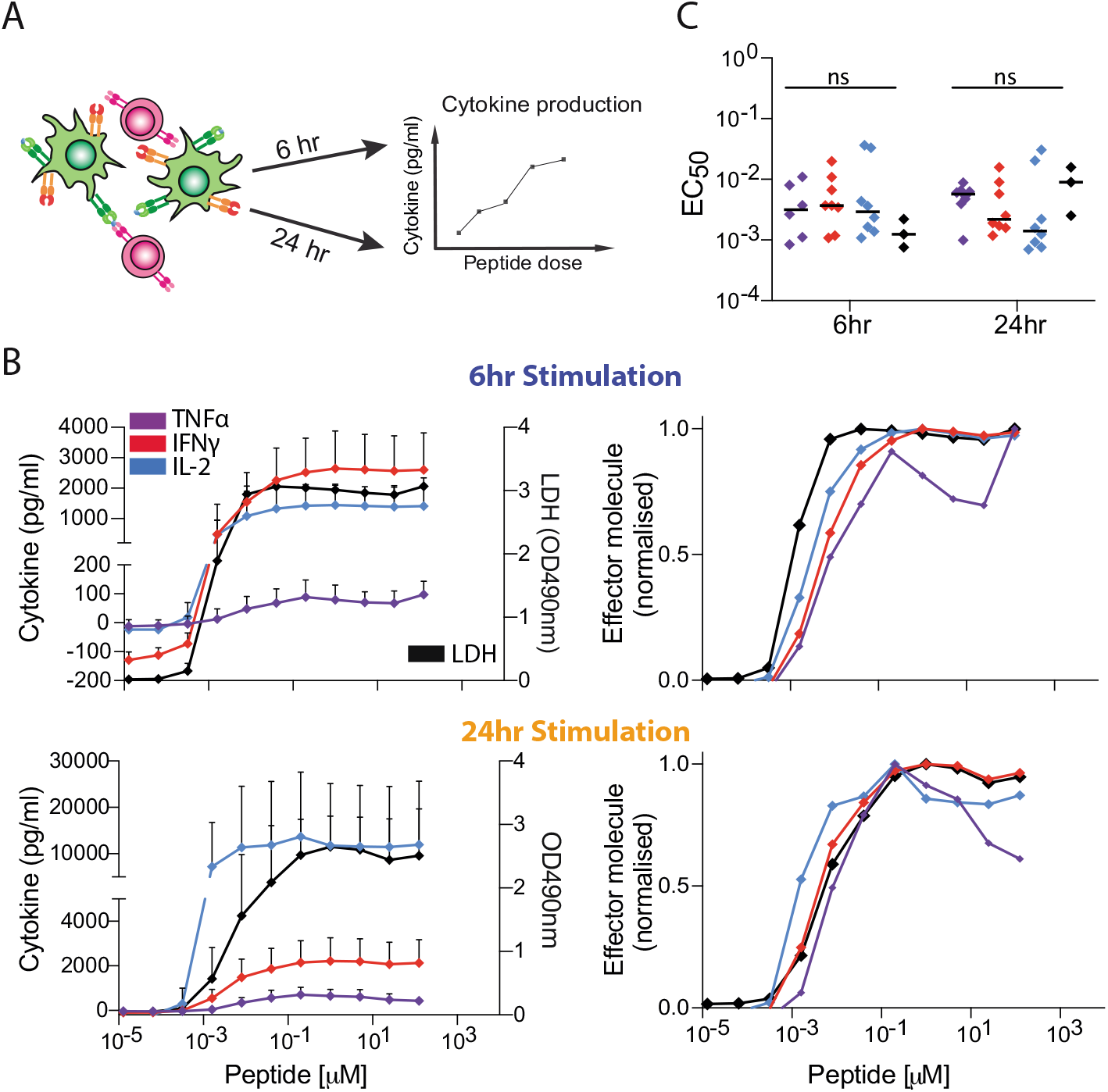
Memory CD8^+^ T cells produce different cytokines and induce killing at a comparable antigen threshold. A) Schematic of experimental assay showing memory CD8^+^ T cells electroporated with the 1G4 TCR (magenta) recognising 9V peptides on MHC (orange) loaded onto monocyte-derived dendritic cells (green) for 6 or 24 hr before cytokine or LDH levels are measured in the supernatant. The 1G4 TCR affinity for the 9V is K_D_ = 7 μM (15). B) Average supernatant TNFα, IFNγ and IL-2 (left Y-axis) or LDH (right Y-axis) as a function of peptide concentration following T cell activation by peptide loaded mature monocyte-derived dendritic cells for 6 hr (top) or 24 hr (bottom) from at least 3 donors. Error bars are SD of mean. C) The EC_50_ for each effector molecule from multiple donors are plotted at 6 and 24 hr (dots are individual donors, horizontal bar is median) showing no significant difference between the different cytokines (ANOVA corrected for multiple tests using Tukey’s test).

## Discussion

Using systematic experiments in a reductionist plate-based system, with precise control of pMHC antigen dose/affinity and co-stimulation through CD28, CD2, and CD27, we found no evidence for different antigen thresholds for different cytokines produced by CD8^+^ T cell blasts. We observed similar results when using memory CD8^+^ T cells stimulated by monocyte-derived antigen presenting cells expressing a host of co-signalling receptor ligands (12).

Co-stimulation by CD2, CD27, and CD28 increased T cell cytokine production in our plate-based reductionist system but were quantitatively distinct (Fig. 3, 4, 5). In all cases, co-stimulation increased the absolute amount of cytokine produced (increased in E_max_) and increased antigen potency (decrease EC_50_). However, the fold-increase in E_max_ and the fold-decrease in EC_50_ were largest for CD2 and not for the more canonical co-stimulation receptor CD28, whereas CD27 exhibited only modest fold-changes. Although CD2 was initially reported to have only subtle role in T cell activation in mice (25, 26), it is increasingly clear that it is important for human T cell activation (21,27,28) and may be particularly important for CD8^+^ T cells that do not express CD28 (29). Although co-stimulation in this reductionist system clearly controlled the antigen threshold for T cell cytokine production, it appeared to do so similarly for different cytokines. Therefore, we found no evidence that a different antigen threshold elicited different effector cytokines in CD8^+^ T cells.

The discrepancy between our results and previous work may be a result of differences in experimental assays and time points. We have measured population-level supernatant cytokine levels whereas previous work relied on intracellular cytokine staining (6, 8, 9), which provides single-cell information but by blocking secretion may affect different cytokines differently. This difference is apparent in a study that directly compared the two methods showing a different threshold for different cytokines when using intracellular staining but not in the supernatant (see Fig. 2 in (30)). A technically more demanding assay based on single-cell cytokine secretion has shown that a single pMHC can induce both TNFα and IL-2 implying that their antigen threshold is comparable (31). In addition, it is now clear that different cytokines exhibit different production kinetics (32, 33) and production depends on continuous TCR/pMHC engagement (34). Therefore, the pMHC degradation rate may introduce apparent differences in thresholds with cytokines having faster production kinetics appearing to have a lower threshold. To control for this, we have used a highly stable variant of the NY-ESO-1 peptide antigen (9C to 9V) and used recombinant pMHC for constant presentation (18).

We highlight that although the antigen threshold for producing different cytokines may be comparable, or indeed identical, there are multiple mechanisms that enable differential regulation. For example, it is clear that there are differences in bulk cytokine production kinetics (32, 33), which may be a result of different mRNA expression and stability (35–37) and sequential production programs (32). Although differences in individual cytokine levels are clearly observed in individual experiments (see representative curves in Fig. 2C, 3C, 4C, 5C), variability across human donors has meant that these differences were not always statistically significant (see E_max_ values in dot plots). Therefore, although the decision to produce cytokine is tightly coupled to a common antigen threshold, the kinetics of production and the absolute amount can be regulated differently.

Our findings support a molecular signalling model whereby a digital signalling switch is rate-limiting for all cytokines with differences in the absolute amount arising as a result of different production kinetics downstream (Fig. 1B). A digital switch has been reported in the TCR signalling pathway (10, 11) and a large number of T cell responses, including cytokine production, have been shown to be digital (31, 38, 39). Therefore, the observation that the induction of different cytokines have a comparable antigen threshold implies that they have comparable TCR signalling thresholds, and it is likely that they share a common rate-limiting switch. As discussed above, different production kinetics can arise from a variety of mechanisms downstream of the rate-limiting switch (e.g. different mRNA stability). If this switch is proximal to the TCR, it would imply that other effector responses share the same antigen threshold as cytokines. In experiments with APCs, we observed a similar antigen threshold between cytokines and a proxy for killing (Fig. 6). This is consistent with the observation that both killing and cytokine production can be observed in response to <5 pMHC per APC (31,40,41).

The systematic analysis of multiple T cell responses we have performed suggests that antigen recognition switches “on” CD8^+^ T cell effector functions implying a common antigen dose threshold and this common threshold depends on antigen affinity. This model is conceptually appealing because the antigen dose and affinity are chance factors that do not necessarily encode any pathogen-specific information that may favour one effector response over another. In this model, pathogen-specific information may be encoded by TCR-extrinsic factors, such as ligands to other co-signalling receptors (42). This is consistent with recent *in-vitro* (43) and *in-vivo* (44) data showing that different antigen doses and affinities produce CD8^+^ T cells with similar response potentials. This conclusion may differ for CD4^+^ T cell differentiation, where antigen dose can selectively induce regulatory T cells, Th1, and Th2 phenotypes (45–47). In summary, we propose that T cell effector responses are maintained by a common critical antigen-dependent decision that can be subsequently regulated by temporal-integration and extrinsic co-signalling receptors that can be response-specific.

## Materials & Methods

### Protein production

pMHCs were refolded in *vitro* from the extracellular residues 1-287 of the HLA-A*02:01 α-chain, β2-microglobulin and NY-ESO-1_157-165_ peptide variants as described previously (17). CHO cell lines permanently expressing the extracellular part of human CD86 (amino acids 6-247) or human CD58 (amino acids 29-213) with a His-tag for purification and a BirA biotinylation site were kindly provided by Simon Davis (University of Oxford, UK). Cells were cultured in GS selection medium and passaged every 3-4 days. After 4-5 passages from thawing a new vial, cells from 2 confluent T175 flasks were transferred into a cell factory and incubated for 5-7 days after which the medium was replaced. The supernatant was harvested after another three weeks, sterile filtered and dialysed over night. The His-tagged protein (CD86 or CD58) was purified on a Nickel-NTA Agarose column.

CD70 (CD27 ligand) expression constructs were a kind gift from Harald Wajant (Wuerzburg, Germany) and contained a Flag-tag for purification and a tenascin-C trimerisation domain. We added a BirA biotinylation site. The TM protein was produced by transient transfection of HEK 293T cells with XtremeGene^TM^ HP Transfection reagent (Roche) according to the manufacturer’s instructions and purified following a published protocol (48), with the exception of the elution step where we used acid elution with 0.1 M glycine-HCl at pH 3.5.

The pMHC or costimulatory ligand was then biotinylated *in vitro* by BirA enzyme according to the manufacturer’s instructions (Avidity) and purified using size-exclusion chromatography with HBS-EP (0.01 M HEPES pH 7.4, 0.15 M NaCl, 3 mM EDTA, 0.005% v/v Tween 20) as flow buffer and stored in aliquots at −80°C.

### Production of lentivirus for transduction

HEK 293T cells were seeded into 6-well plates before transfection to achieve 50-80% confluency on the day of transfection. Cells were co-transfected with the respective third-TM generation lentiviral transfer vectors and packaging plasmids using Roche XtremeGene^TM^ 9 (0.8μg lentiviral expression plasmid, 0.95 μg pRSV-rev, 0.37 μg pVSV-G, 0.95 μg pGAG). The supernatant was harvested and filtered through a 0.45 μm cellulose acetate filter 24-36h later. The 1G4 TCR used for this project was initially isolated from a melanoma patient (13). The affinity maturation to the c58c61 variant used herein (referred to as 1G4^Hi^) was carried out by Adaptimmune Ltd (14). The TCR in this study has been used in a standard third-generation lentiviral vector with the EF1 α promoter.

### T cell isolation and culture

Up to 50 ml peripheral blood were collected by a trained phlebotomist from healthy volunteer donors after informed consent had been taken. This project has been approved by the Medical Sciences Inter-Divisional Research Ethics Committee of the University of Oxford (R51997/RE001) and all samples were anonymised in compliance with the Data Protection Act. Alternatively, leukocyte cones were purchased from National Health Services Blood and Transplant service. Only HLA-A2^-^ peripheral blood or leukocyte cones were used due to the cross-reactivity of the high-affinity receptors used in this project which leads to fratricide of HLA-A2^+^ T cells (30). CD8^+^ T cells were isolated directly from blood using the CD8^+^ T Cell Enrichment Cocktail (StemCell Technologies) and density gradient centrifugation according to the kit’s instructions. The isolated CD8^+^ T cells were washed and resuspended at a concentration of 1 × 10^6^ cells per ml in completely reconstituted RPMI supplemented with 50 units/ml IL-2 and 1 × 10^6^ CD3/CD28-coated Human T-Activator Dynabeads (Life Technologies) per ml. The next day, 1 × 10^6^ T cells were transduced with the 2.5 ml virus-containing supernatant from one well of HEK cells supplemented with 50 units/ml of IL-2. The medium was replaced with fresh medium containing 50 units/ml IL-2 every 2–3 d. CD3/CD28-coated beads were removed on day 5 after lentiviral transduction and the cells were used for experiments on days 10-14.TCR expression was assessed by staining with NY-ESO 9V PE-conjugated tetramer (in-house produced using refolded HLA*A0201 with NY-ESO 9V and Streptavidin-PE [AbD Serotec or Biolegend]) using flow cytometry.

### T cell stimulation

T cells were stimulated with titrations of plate-immobilised pMHC ligands with or without co-immobilised ligands for accessory receptors. Ligands were diluted to the working concentrations in sterile PBS. 50 μl serially two-fold diluted pMHC were added to each well of high-binding capacity of Streptavidin-coated 96-well plates (15500, Thermo Fisher). After a minimum 45 min incubation at room temperature, the plates were washed once with sterile PBS. Where accessory receptor ligands were used, those were similarly diluted and added to the plate for a second incubation of 45-90 min. After washing the stimulation plate with PBS, 7.5 × 10^4^ T cells were added in 200 μl complete RPMI without IL-2 to each stimulation condition. The plates were spun at 50-200 x g for 2 min to settle down the cells and then incubated at 37 °C with 5 % CO_2_ for 8 hr.

### T cells-Antigen presenting cells stimulation assay

The assay was performed as previously described (12). Briefly, memory CD8^+^ T cells were isolated from anonymised leukapheresis products obtained from the NHS at Oxford University Hospitals by (REC 11/H0711/7), using memory CD8^+^ isolation kit (StemCell Technologies). T cells were harvested and washed three times with Opti-MEM (LifeTechnologies). The cells were resuspended at 25 x 10^6^/ml and 2.5-5 x 10^6^ cells were mixed with the desired mRNA products and aliquoted into 100-200 μl per electroporation cuvette (Cuvette Plus, 2 mm gap, BTX). For each 10^6^ cells CD8 T cells; 5 μg of each TCRα, TCRβ and CD3ζ RNA was used. Cells were electroporated at 300V, 2ms in an ECM 830 Square Wave Electroporation System (BTX). Cells were used after 24 h. Monocytes were enriched from the same leukapheresis products using RosetteSep kits (Stemcell Technologies), and were then cultured at 1-2 x 10^6^/ml in 12-well plates with differentiation media containing 50 ng/ml Interleukin 4 (Peprotech) and 100 ng/ml granulocyte-monocyte colony stimulating factor (GM-CSF, Immunotools) for 24 hr. For maturation the following cytokines were added for an additional 24 hr, 1 μM Prostaglandin E2 (PGE2, Sigma), 10 ng/ml Interleukin 1 beta (Biotechne), 50 ng/ml Tumour Necrosis Factor alpha (Peprotech), and 20 ng/ml Interferon gamma (Bio-techne). Monocyte-derived dendritic cells were loaded with peptide for 90 min at 37°C. T cells and dendritic cells were mixed at 1:1 ratio, 50,000 cells each, and incubated for 6 or 24 h before supernatant was collected and analysed.

### ELISA

Supernatants from stimulation experiments were used fresh. Cytokine concentrations were measured by TM ELISAs according to the manufacturer’s instructions in Nunc MaxiSorp^TM^ flat-bottom plates (Invitrogen) using Uncoated ELISA Kits (Invitrogen) for TNF-α, IFN-γ, and IL-2. LDH Cytotoxicity Detection kit (Takra Bio) was used as per manufacturer instructions to detect cell killing.

### Data analysis

A smoothing function with 10,000 data points was fitted to the experimental data to empirically extract the maximum amount of cytokine produced across different pMHC concentrations (E_max_) and the pMHC concentration producing a half-maximal response (EC_50_) for each dose-response curve. The difference of two sigmoidal dose-response curves was used as a smoothing function to capture the bell-shaped dose-response curve frequently observed (17) and it was fitted to the experimental data using the *lsqcurvefit* function in Matlab (Mathworks, MA).

Experiments with human donors are often highly variable and we found large quantitative differences in the range of cytokine production between experimental repeats. Before averaging across different donors for panels B in each figure, the E_max_ and EC_50_ were thus, for each donor, normalised to the mean E_max_ and mean EC_50_, respectively, for all the cytokines across all pMHC affinities (Fig. 2, Fig. S2) or all doses of the co-stimulatory ligand (Fig. 3, Fig. 4, Fig. 5, Fig. S3, Fig. S4, Fig. S5). The normalised E_max_ and EC_50_ were then averaged across all human donors and they are presented in panels B of each figure as a fold-change relative to the respective metric for IL-2 in response to the highest-affinity pMHC in Fig. 2 or the experimental control with pMHC alone in Fig. 3–5 and Fig. S3–S5 to allow for a more intuitive comparison of cytokines.

Similarly, before averaging across different donors for dotplots in each figure, the E_max_ and EC_50_ were, for each donor, normalised to the mean E_max_ and mean EC_50_, respectively, for any given pMHC affinity (Fig. 2) or dose of the co-stimulatory ligand (Fig. 3–5, Fig. S4–S5). The normalised E_max_ and EC_50_ were then averaged across all human donors and they are again presented in Fig. S2–S5 as a fold-change relative to the respective metric for IL-2 to more directly identify potential differences in E_max_ and EC_50_ between cytokines at each pMHC affinity and concentration of co-stimulatory ligand.

### Statistical analysis

A nonparametric Spearman correlation test was used to identify whether pMHC affinity (Fig. 2) or the concentration of co-stimulatory ligands (Fig. 3–5, Fig. S4–S5) correlated with the E_max_ and EC_50_ of each cytokine response (Panels B in all figures). Ordinary repeated-measures two-way ANOVA corrected for multiple comparisons by Tukey’s test was performed on experimental data to determine whether differences in E_max_ and EC_50_ at each pMHC affinity (Fig. 2) or concentration of co-stimulatory ligand (Fig. 3–5, Fig. S3–S5) were statistically significant between cytokines. GraphPad Prism was used for all statistical analyses.

## Acknowledgements

We thank Simon J. Davis for providing CD86 and CD58 expression plasmids, Harald Wajant for providing CD70 expression plasmids, Adaptimmune Ltd for providing the c58c61 TCR. We thanks all members of Omer Dushek lab, and Michael L. Dustin for feedback on experimental protocols and discussion of the manuscript. We thank P. Anton van der Merwe and Marion H. Brown for a critical reading of the manuscript.

## Author contributions

EAS, NT, PK, JN, JP and MK performed experiments; EAS and NT analysed data; EAS, NT and OD designed the research and wrote the paper; EAS and NT contributed equally. All authors discussed the results and commented on the paper.

## Supplementary Information

### Supplementary Figures

**Figure S1:**
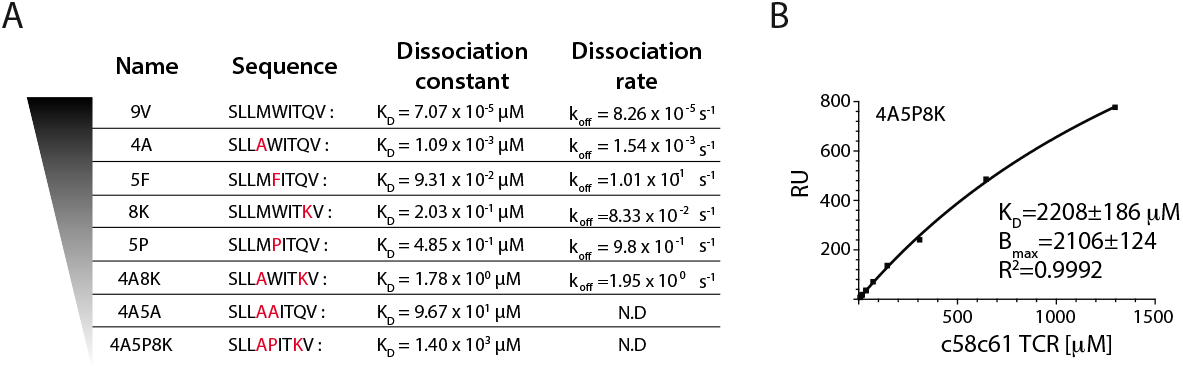
Binding parameters for the panel of pMHC used in the study. A) The dissociation constant and dissociation rate for the 8 pMHC used meausred by surface plasmon resonance (SPR) at 37 degrees. All parameters except for 4A5A and 4A5P8K were reported previously (17). B) SPR experiments were repeated for the two lowest affinity pMHC at 37 degrees as previously described (17) with higher TCR concentrations to more accurately determine the dissociation constant. Representative binding response at steady-state over the TCR concentration is shown for 4A5P8K. The average of at least 3 independent experiments were used to calculate the KD values in panel A.

**Figure S2:**
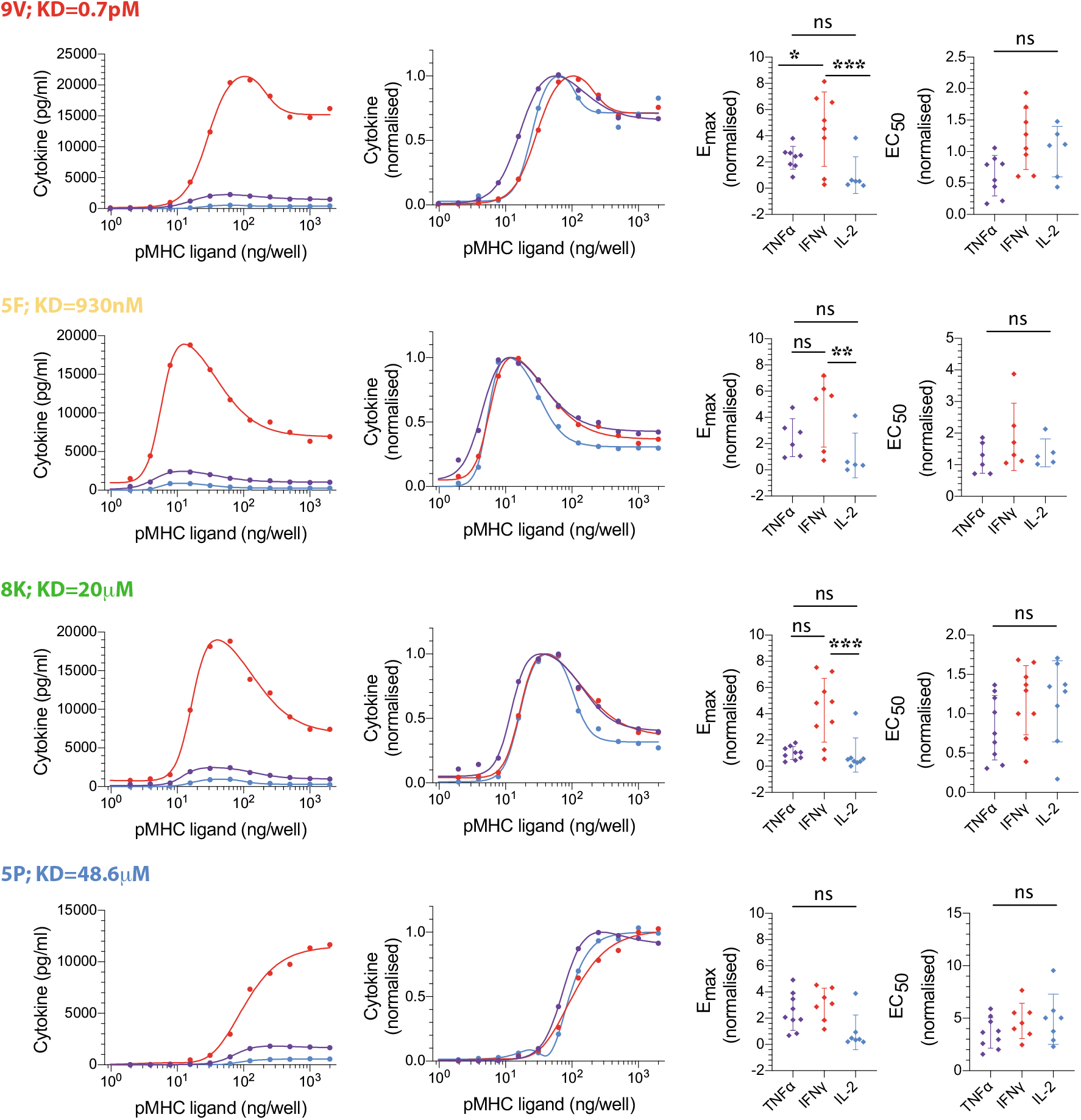
Changes to the pMHC affinity does not introduce differences in EC_50_ for each cytokine. Direct comparison of normalised E_max_ (left dotplot) and EC_50_ (right dotplot) for TNFα, IFNγ and IL-2 for different affinity pMHC (rows) showing significant but < 2-fold differences between IFNγ and IL-2 on the one hand and TNFα on the other hand. Error bars are SD of mean. Normalisation of experimental data is described in Materials and Methods. ANOVA corrected for multiple comparisons by Tukey’s test (*; p=0.0383, **; p=0.006, ***; pi0.001).

**Figure S3:**
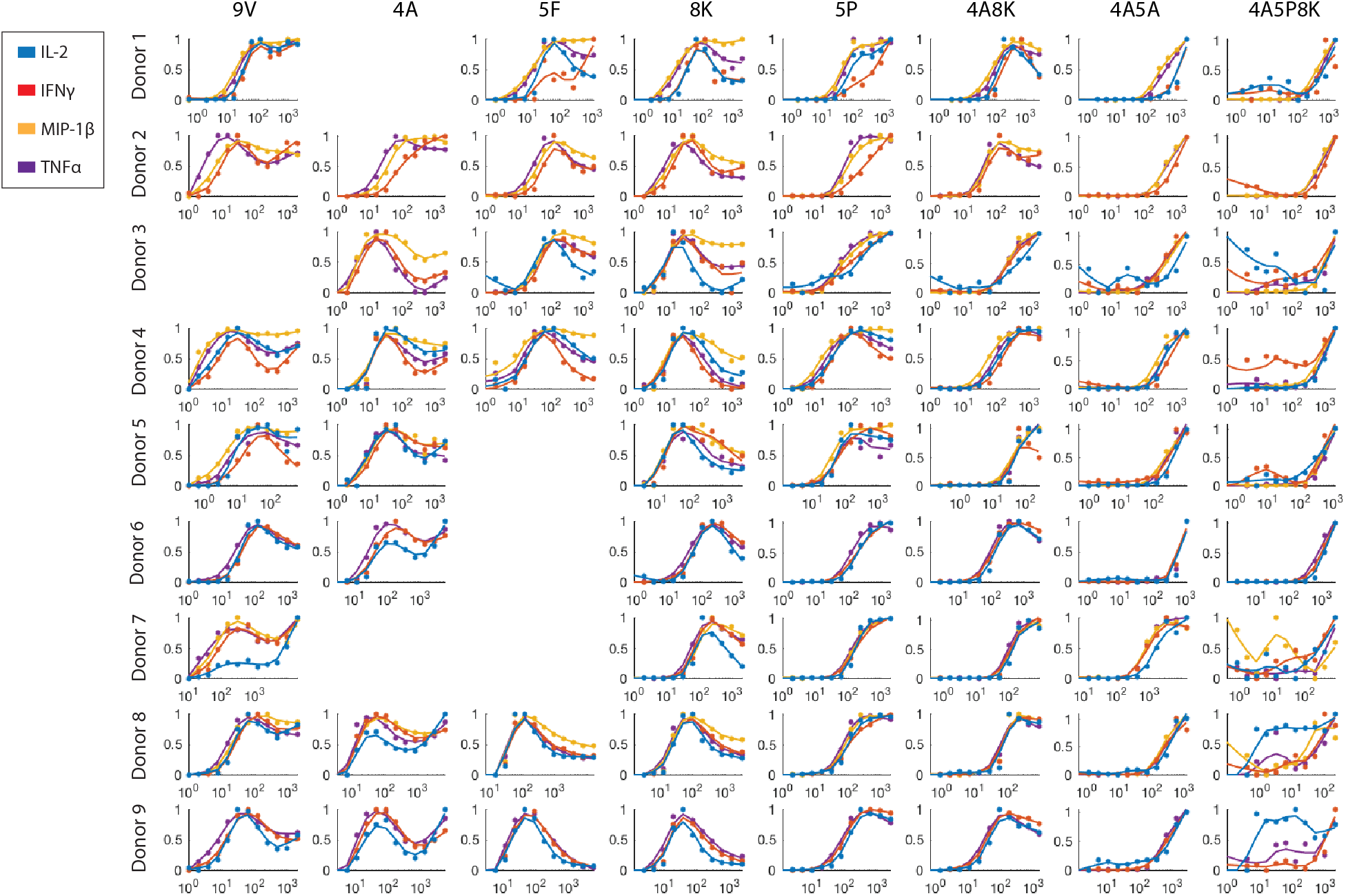
Overlay of cytokine dose-response curves highlights that different cytokines have a comparable antigen dose threshold and this conclusion holds for different antigen affinity. The dose-response curves for TNFα, IFNγ, MIP1β and IL-2 were normalised by their respective E_max_ value for different affinity pMHCs (columns) and individual donors (rows). Although the two lowest affinity pMHCs (4A5A, 4A5P8K) were omitted from the quantitative analysis because accurate estimates of EC_50_ were not possible, representative curves highlight that they also share a common antigen threshold. Additionally, MIP1β exhibited a shared antigen threshold but was omitted from the main text because it was not measured in all experimental conditions. Normalisation of experimental data is described in Materials and Methods.

**Figure S4:**
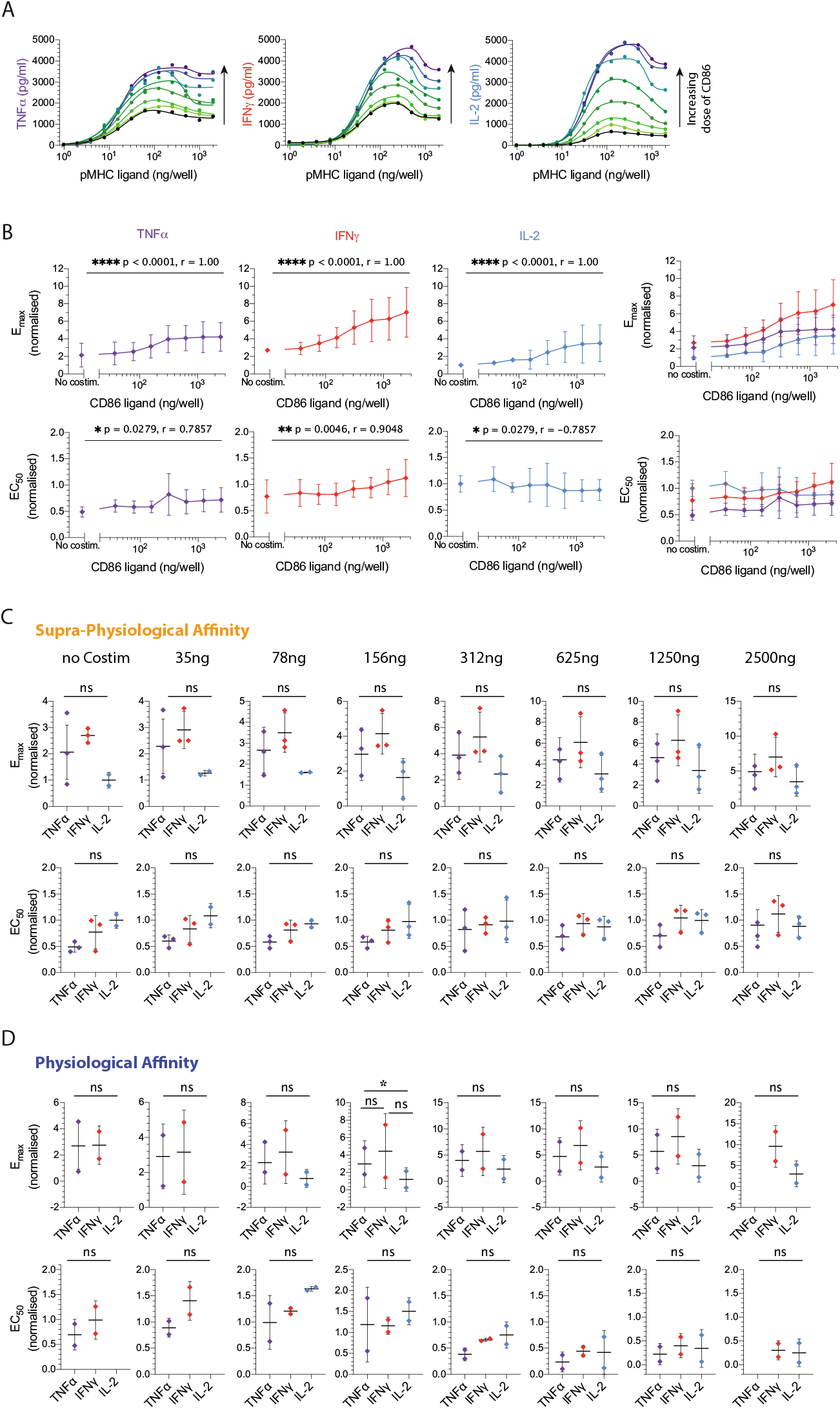
CD28 co-stimulation does not alter the threshold for cytokine secretion. A) Representative dataset showing secretion of TNFα, IFNγ and IL-2 as a function of supra-physiological affinity pMHC dose (4A) when T cells were costimulated with different doses of CD86 ligand (colours). B) Top row: Normalised E_max_ of cytokine produced as a function of CD86 dose confirms that co-stimulation amplifies cytokine secretion for individual cytokines or an overlay of all three (right panel). Bottom row: CD86 co-stimulation has a minor impact on the EC_50_ of cytokine secretion in response to the supra-physiological-affinity pMHC mutant. C-D) Direct comparison of normalised E_max_ (top) and EC_50_ (bottom) for TNFα, IFNγ and IL-2 for different doses of CD86 ligand (columns) showing no significant and < 2fold differences between the three cytokines independent of CD86 co-stimulation for (C) supra-physiological, from the plots in B and (D) physiological-affinity pMHC (D) from plots in Fig. 3. Error bars are SD of mean. Normalisation of experimental data is described in Materials and Methods. Solid lines in representative datasets are the fits used to extract E_max_ and EC_50_. ANOVA corrected for multiple comparisons by Tukey’s test (*; p=0.0266).

**Figure S5:**
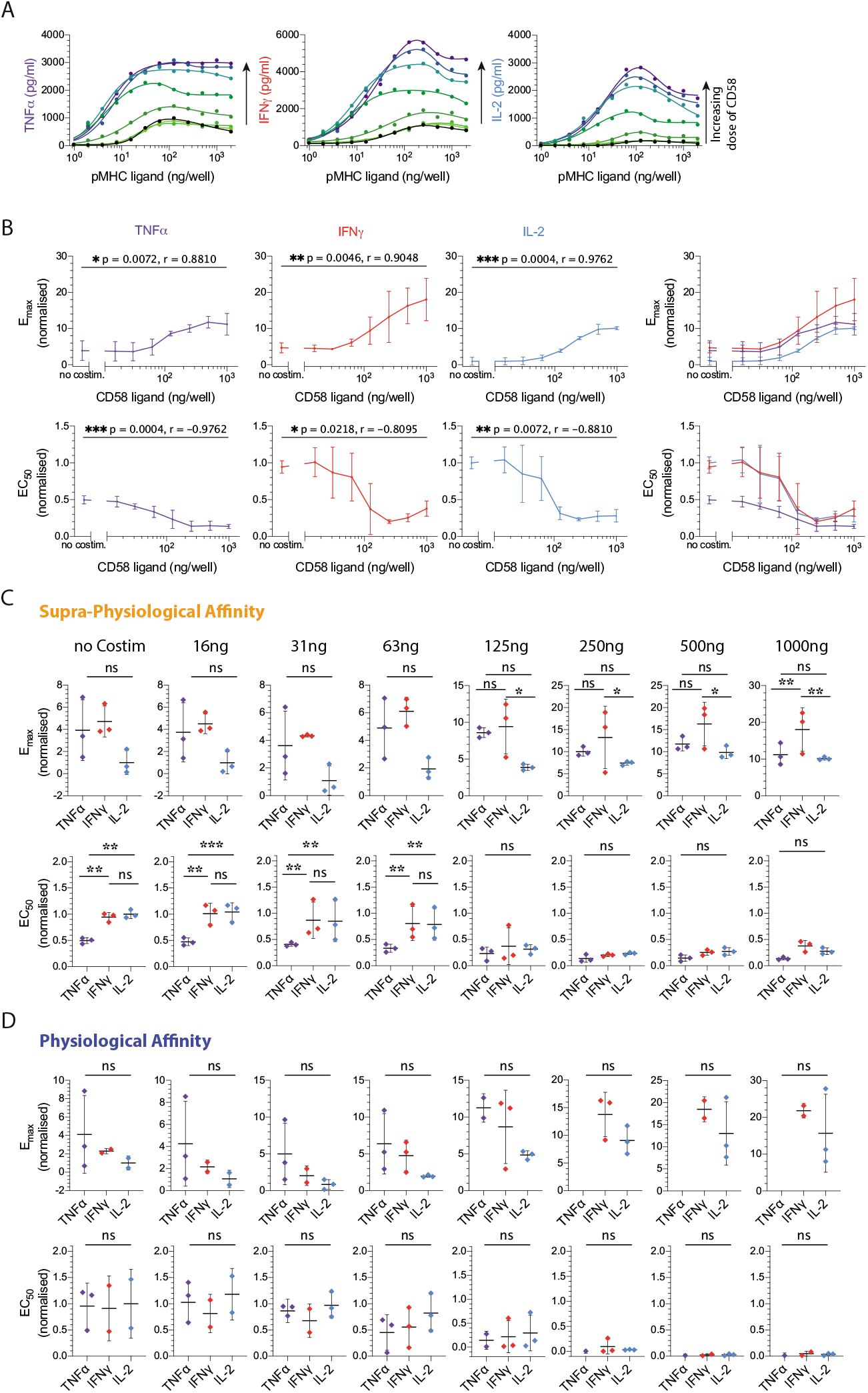
CD2 co-stimulation does not alter the threshold for cytokine secretion. A) Representative dataset showing secretion of TNFα, IFNγ and IL-2 as a function of supra-physiological affinity pMHC (4A) dose when T cells were costimulated with different doses of CD58 (colours). B) Top row: Normalised E_max_ of cytokine produced as a function of CD58 dose confirms that co-stimulation amplifies cytokine secretion for individual cytokines or an overlay of all three (right panel). Bottom row: CD58 co-stimulation has a profound impact on the EC_50_ of cytokine secretion in response to the supra-physiological-affinity pMHC mutant. C-D) Direct comparison of normalised E_max_ (top) and EC_50_ (bottom) for TNFα, IFNγ and IL-2 for different doses of CD58 (columns) showing no significant and < 2-fold differences between the three cytokines independent of CD58 co-stimulation for (C) supra-physiological, from the plots in B and (D) physiological-affinity pMHC (D) from plots in Fig. 4. Error bars are SD of mean. Normalisation of experimental data is described in Materials and Methods. Solid lines in representative datasets are the fits used to extract E_max_ and EC_50_. ANOVA corrected for multiple comparisons by Tukey’s test (**; p=0.005, ***; p=0.0007).

## References

1. Smith-Garvin JE, Koretzky GA, Jordan MS (2009) T cell activation. Annual review of immunology 27:591–619.

2. Paul WE, Seder RA (1994) Lymphocyte responses and cytokines. Cell 76:241–251.

3. Moticka EJ (2015) A historical perspective on evidence-based immunology (Newnes).

4. Pennock ND, et al. (2013) T cell responses: naive to memory and everything in between. Advances in physiology education 37:273–283.

5. Valitutti S, Müller S, Dessing M, Lanzavecchia A (1996) Different responses are elicited in cytotoxic t lymphocytes by different levels of t cell receptor occupancy. The Journal of experimental medicine 183:1917–1921.

6. van den Berg HA, et al. (2013) Cellular-level versus receptor-level response threshold hierarchies in t-cell activation. Frontiers in immunology 4:250.

7. Price DA, et al. (1998) Antigen–specific release of β-chemokines by anti-hiv-1 cytotoxic t lymphocytes. Current biology 8:355–358.

8. Itoh Y, Germain RN (1997) Single cell analysis reveals regulated hierarchical t cell antigen receptor signaling thresholds and intraclonal heterogeneity for individual cytokine responses of cd4+ t cells. The Journal of experimental medicine 186:757–766.

9. Hemmer B, Stefanova I, Vergelli M, Germain RN, Martin R (1998) Relationships among tcr ligand potency, thresholds for effector function elicitation, and the quality of early signaling events in human t cells. The Journal of Immunology 160:5807–5814.

10. Das J, et al. (2009) Digital signaling and hysteresis characterize ras activation in lymphoid cells. Cell 136:337–51.

11. Altan-Bonnet G, Germain RN (2005) Modeling T cell antigen discrimination based on feedback control of digital ERK responses. PLoS biology 3:e356.

12. Abu-Shah E, et al. (2019) A tissue-like platform for studying engineered quiescent human t-cells’ interactions with dendritic cells. eLife 8.

13. Chen JL, et al. (2000) Identification of ny-eso-1 peptide analogues capable of improved stimulation of tumor-reactive ctl. The Journal of Immunology 165:948–955.

14. Li Y, et al. (2005) Directed evolution of human t-cell receptors with picomolar affinities by phage display. Nature biotechnology 23:349–354.

15. Aleksic M, et al. (2010) Dependence of T cell antigen recognition on T cell receptor-peptide MHC confinement time. Immunity 32:163–74.

16. Dushek O, et al. (2011) Antigen potency and maximal efficacy reveal a mechanism of efficient T cell activation. Science Signaling 4:ra39.

17. Lever M, et al. (2016) Architecture of a minimal signaling pathway explains the t-cell response to a 1 millionfold alitariation in antigen affinity and dose. Proceedings of the National Academy of Sciences 113:E6630–E6638.

18. Trendel NC, Kruger P, Nguyen J, Gaglione S, Dushek O (2019) Perfect adaptation of cd8+ t cell responses to constant antigen input over a wide range of affinity is overcome by costimulation. BioRxiv p 535385.

19. Harding FA, McArthur JG, Gross JA, Raulet DH, Allison JP (1992) Cd28-mediated signalling co-stimulates murine t cells and prevents induction of anergy in t-cell clones. Nature 356:607–609.

20. Springer TA (1990) Adhesion receptors of the immune system. Nature 346:425–434.

21. Demetriou P, et al. (2019) Cd2 expression acts as a quantitative checkpoint for immunological synapse structure and t-cell activation. bioRxiv p 589440.

22. Kaizuka Y, Douglass AD, Vardhana S, Dustin ML, Vale RD (2009) The coreceptor cd2 uses plasma membrane microdomains to transduce signals in t cells. Journal of Cell Biology 185:521–534.

23. Ward-Kavanagh LK, Lin WW, Šedý JR, Ware CF (2016) The TNF Receptor Superfamily in Co-stimulating and Co-inhibitory Responses. Immunity 44:1005–1019.

24. Legat A, Speiser DE, Pircher H, Zehn D, Fuertes Marraco SA (2013) Inhibitory receptor expression depends more dominantly on differentiation and activation than ‘‘exhaustion” of human CD8 T cells. Frontiers in Immunology 4:1–15.

25. van der Merwe Pa (1999) A subtle role for CD2 in T cell antigen recognition. The Journal of Experimental Medicine 190:1371–1374.

26. Bachmann MF, Barner M, Kopf M (1999) CD2 sets quantitative thresholds in T cell activation. Journal of Experimental Medicine 190:1383–1392.

27. Patel SJ, et al. (2017) Identification of essential genes for cancer immunotherapy. Nature 548:537–542.

28. Wang ECY, et al. (2018) Suppression of costimulation by human cytomegalovirus promotes evasion of cellular immune defenses. Proceedings of the National Academy of Sciences 115:4998–5003.

29. Leitner J, Herndler-Brandstetter D, Zlabinger GJ, Grubeck-Loebenstein B, Steinberger P (2015) CD58/CD2 Is the Primary Costimulatory Pathway in Human CD28 — CD8 + T Cells. The Journal of Immunology 195:477–487.

30. Tan M, et al. (2015) T cell receptor binding affinity governs the functional profile of cancer-specific cd8+ t cells. Clinical & Experimental Immunology 180:255–270.

31. Huang J, et al. (2013) A single peptide-major histocompatibility complex ligand triggers digital cytokine secretion in cd4+ t cells. Immunity 39:846–857.

32. Han Q, et al. (2012) Polyfunctional responses by human T cells result from sequential release of cytokines. Proceedings of the National Academy of Sciences 109:1607–1612.

33. Salerno F, Paolini NA, Stark R, von Lindern M, Wolkers MC (2017) Distinct PKC-mediated posttranscriptional events set cytokine production kinetics in CD8 <sup>+</sup> T cells. Proceedings of the National Academy of Sciences 114:201704227.

34. Huppa JB, Gleimer M, Sumen C, Davis MM (2003) Continuous T cell receptor signaling required for synapse maintenance and full effector potential. Nature immunology 4:749–55.

35. Seko Y, Cole S, Kasprzak W, Shapiro BA, Ragheb JA (2006) The role of cytokine mrna stability in the pathogenesis of autoimmune disease. Autoimmunity reviews 5:299–305.

36. Nicolet BP, Guislain A, Wolkers MC (2017) Combined single-cell measurement of cytokine mrna and protein identifies t cells with persistent effector function. The Journal of Immunology 198:962–970.

37. Friedman RL, Manly SP, McMahon M, Kerr IM, Stark GR (1984) Transcriptional and posttranscriptional regulation of interferon-induced gene expression in human cells. Cell 38:745–755.

38. Podtschaske M, et al. (2007) Digital NFATc2 activation per cell transforms graded T cell receptor activation into an all-or-none IL-2 expression. PLoS ONE 2.

39. Mayya V, Dustin ML (2016) What Scales the T Cell Response? Trends in Immunology 37:513–522.

40. Purbhoo MA, Irvine DJ, Huppa JB, Davis MM (2004) T cell killing does not require the formation of a stable mature immunological synapse. Nature Immunology 5:524–530.

41. Siller-Farfán JA, Dushek O (2018) Molecular mechanisms of T cell sensitivity to antigen. Immunological Reviews 285:194–205.

42. Chen L, Flies DB (2013) Molecular mechanisms of T cell co-stimulation and co-inhibition. Nature Reviews Immunology 13:227–242.

43. Richard AC, et al. (2018) T cell cytolytic capacity is independent of initial stimulation strength. Nature immunology 19:849–858.

44. Ozga AJ, et al. (2016) pmhc affinity controls duration of cd8+ t cell-dc interactions and imprints timing of effector differentiation versus expansion. Journal of Experimental Medicine 213:2811–2829.

45. Panhuys NV, Klauschen F, Germain R (2014) T-Cell-Receptor-Dependent Signal Intensity Dominantly Controls CD4+ T Cell Polarization In Vivo. Immunity pp 1–12.

46. Turner MS, Kane LP, Morel Pa (2009) Dominant role of antigen dose in CD4+ Foxp3+ regulatory T cell induction and expansion. The Journal of Immunology 183:4895–4903.

47. Gottschalk Ra, Corse E, Allison JP (2010) TCR ligand density and affinity determine peripheral induction of Foxp3 in vivo. The Journal of experimental medicine 207:1701–11.

48. Wyzgol A, et al. (2009) Trimer stabilization, oligomerization, and antibody-mediated cell surface immobilization improve the activity of soluble trimers of cd27l, cd40l, 41bbl, and glucocorticoid-induced tnf receptor ligand. The Journal of Immunology 183:1851–1861.

